# Microvascular architecture and dynamics of the choroid plexus brain barrier

**DOI:** 10.64898/2026.03.16.711715

**Authors:** Samantha Kuszynski, Ian Junker, Shristi Shrestha, Alexander Brand, Paige Pfotenhauer, Oleg Kovtun, Ryan Moran, Chloe Koo, Carson Oakes, José Maldonado, Jean-Phillipe Cartailler, Alex Tiriac, Neil Dani

**Author notes:** Co-first authors (Samantha Kuszynski, Ian Junker).

## Abstract

The choroid plexus is a specialized blood-cerebrospinal fluid barrier that supports cerebrospinal fluid production, immune surveillance, and molecular exchange between the circulation and the central nervous system, yet its vascular bed remains poorly understood. Here, we combine whole-tissue clearing and light-sheet imaging, single-nucleus transcriptomic reanalysis, and live calcium imaging of intact murine explants to define the structural and functional organization of choroid plexus endothelial networks across development. Three-dimensional imaging and reconstruction highlights a dense, epithelial-ensheathed vascular plexus that is continuous with the broader cerebrovasculature and organized into anatomically distinct inflow and ventricular margin regions. Transcriptomic analysis identified developmentally stratified endothelial subtypes, with embryonic populations enriched for proliferative, motor, and membrane remodeling programs and adult and aged populations enriched for adhesion, extracellular matrix, transport, and mechanosensory pathways. Endothelial subsets across stages expressed genes linked to flow sensing and calcium-dependent mechanotransduction, including *Piezo1*, *Piezo2*, and *Trpv4*. Consistent with these signatures, intact explants exhibited spontaneous, spatially graded calcium oscillations, and pharmacologic activation by Piezo1, triggered robust network-wide calcium responses in embryonic and adult tissue with distinct temporal dynamics. Piezo1 activation also promoted stabilization of PECAM1-associated endothelial adhesion under *ex vivo* flow conditions. Together, these findings establish the choroid plexus endothelium as a structurally specialized, developmentally dynamic, and mechanosensitive vascular network and provide a framework for investigating endothelial contributions to blood-cerebrospinal fluid barrier function in health and disease.

## INTRODUCTION

The choroid plexus, located within the ventricles of the brain, carries out functions vital to brain homeostasis and health throughout life.^1,2^ These include cerebrospinal fluid (CSF) production, ligand secretion, immune signaling, and regulation of factor exchange between the blood and CSF.^3–9^ Next generation sequencing technologies, single-cell transcriptomic studies, and imaging advances have transformed our understanding of choroid plexus gene expression profiles, cellular diversity, and structural foundations.^10–13^ To date, the bulk of choroid plexus research has focused on epithelial and immune cells, while the endothelial cells that form the vascular bed of the tissue remain overlooked. Choroid plexus endothelial cells form a dense vascular plexus and supply water, nutrients, ions, and signaling factors to the choroid plexus and ultimately the CSF. While directly connected to the rest of the brain vasculature, most basic and clinical research has focused on the parenchymal vessels that form the blood brain barrier (BBB).^14–16^ Choroid plexus endothelial cells collaborate with a specialized set of epithelial, mural, neural, glial, and immune cell types to form a functionally distinct blood-CSF barrier.^10,17,18^ Choroid plexus endothelial cells also harbor fenestrae, which are naturally occurring apertures that are better appreciated in circumventricular organs, ocular vascular beds, and peripheral organs such as the kidneys, liver, and pancreas.^20,22,24^ Mechanisms that dynamically regulate permeability of choroid plexus endothelial cells are only beginning to be understood in models of disease, which underscores the need for deeper investigation of fundamental vascular properties that regulate molecular and cellular traffic into and out of the brain.^19^ Research has been limited due to incomplete knowledge and a lack of tools to investigate endothelial cell contributions to choroid plexus functions. Here we present new *in situ* imaging data of the intact choroid plexus vascular bed, their transcriptional diversity, and live imaging strategies to test their function in real time across developing and adult murine brains.

## RESULTS

### Choroid plexus endothelial cells form a plexus that is connected to the cerebrovasculature

The choroid plexus is a highly vascularized blood-CSF interface that is morphologically distinct and anatomically partitioned from brain parenchyma. To visualize *in situ* organization and cell morphology of the choroid plexus vascular bed, we used double fluorescent mT/mG reporter mice to distinguish endothelial cells from non-endothelial cell types.^21^ Crossing reporter mice with an endothelial cell driver (Tek-Cre) enabled Cre-mediated endothelial expression of membrane-targeted enhanced green fluorescence protein (EGFP) while non-endothelial cells remained positive for membrane-targeted tandem dimer Tomato (tdTomato). We harvested and processed brains for imaging from adult mice (postnatal day (P) 56) using the SHIELD protocol followed by active de-lipidation.^28^ Optically clear mouse brains were imaged in three dimensions (3D) using a single-plane illumination light sheet microscope (SmartSPIM, LifeCanvas Technologies, see Methods) followed by stitching and volumetric renderings (Imaris, Oxford Instruments, see Methods) (**Fig. 1A**). To visualize global vascular patterning and local morphology, we pseudocolored endothelial cells (EGFP+) orange and non-endothelial cells (tdTomato+) gray (**Fig. 1B, 1C**). Axial projections of forebrain ventricular spaces showed compartmentalized vascular plexus (orange) enveloped in an epithelial monolayer (gray), consistent with earlier histological studies (**Fig. 1D**). Choroid plexus vessels appeared densely organized with tortuous loops creating a glomerulus-like architecture reminiscent of glandular tissue.^23^ To visualize the relationship with parenchymal vessels, we projected 3D volume stacks of coronal brain sections containing the lateral ventricle, which revealed robust vascular projections to the choroid plexus that branch into capillary loops (**Fig. 1E**). Infiltrating vessels enter through ventral portions of the lateral ventricle choroid plexus and project towards the free margin, or ventricular-facing edge of the tissue (**Fig. 1F-H, S1A**). Surface rendering of the free margin edge shows tufted morphology of epithelial cells that overlay the complex network of underlying endothelial cells that form a plexus (**Fig. 1I, S1B-E**). Quantifying the area of ventricular and stalk portions of the choroid plexus tissue in sequential coronal sections along the rostrocaudal axis of the lateral ventricle, confirmed higher three-dimensional epithelial and vascular complexity in dorsal segments containing the free margin compared to ventral planar portions (**Fig. 1J, 1K, S1F**). Similar space-filling morphology of epithelia-enveloped vasculature was found in choroid plexus explants and coronal sections across all three ventricles (**Fig.S1H-J**).

**Figure 1.**
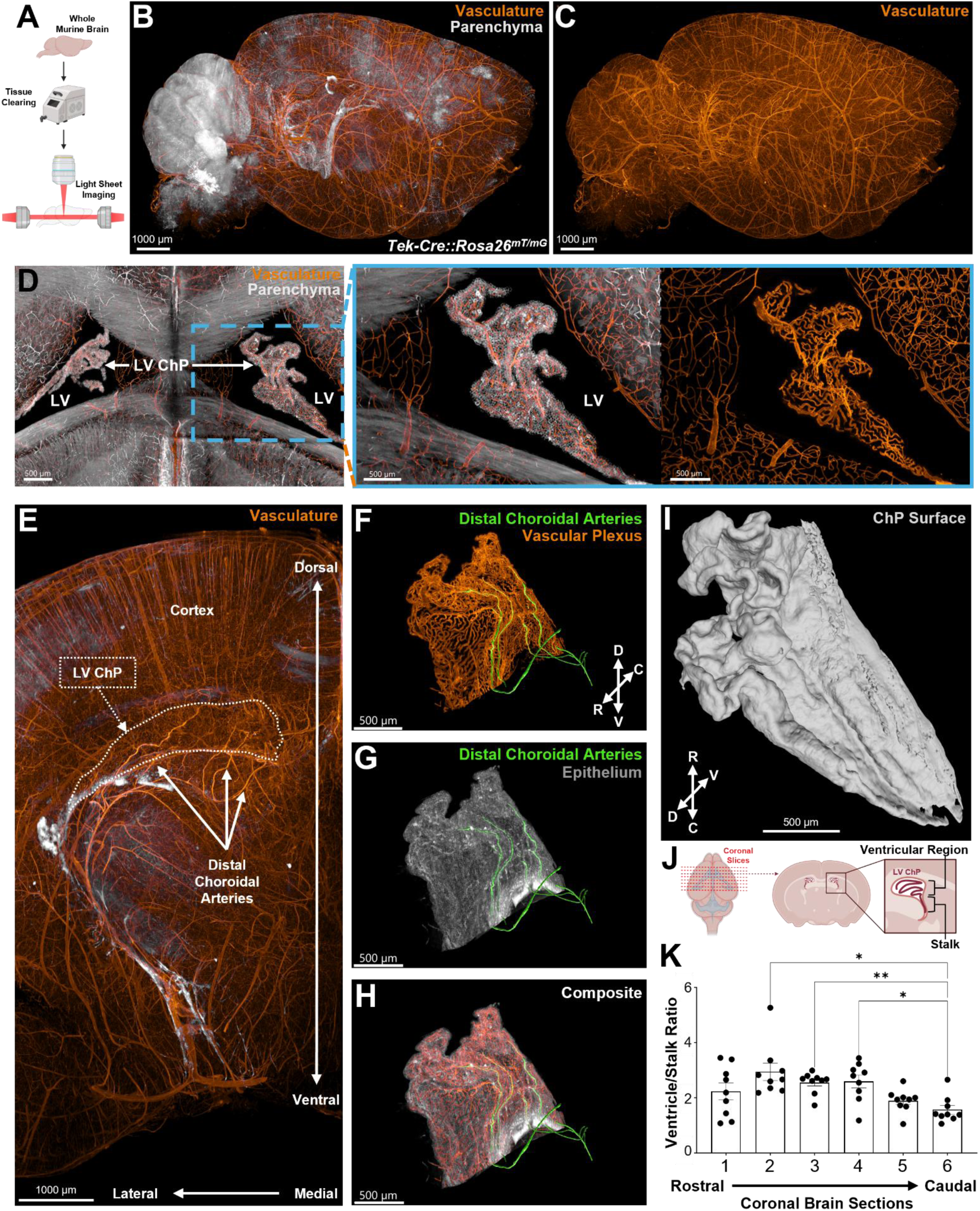
Choroid plexus structure and ventricular organization. (A) Tissue-clearing workflow. (B) Light sheet image of whole murine brain from *Tek-Cre::Rosa26^mT/mG^* reporter mouse. Endothelial cells (mG) and non-endothelial cells (mT) are pseudocolored orange and gray, respectively. (C) Sagittal view of parenchymal vasculature. (D) Axial view of lateral ventricles with zoomed-in view of parenchyma, choroid plexus, and vasculature. (E) Coronal maximum intensity projection (2.5 mm) of a brain hemi-segment shows branching patterns and distal choroidal vessels infiltrating the superior horn of the lateral ventricle choroid plexus. (F-H) Imaris-rendering of segmented lateral ventricular choroid plexus superior horn shows brain-derived infiltrating vessels (green) integrated with choroid plexus vasculature (orange) enveloped by epithelium (gray). (I) 3D model of the superior horn of the lateral ventricle choroid plexus illustrating tufted superstructure along free margin. (J) Quantitation workflow of ventricular and stalk regions of lateral ventricle choroid plexus. (K) Bar graph showing the ration of ventricular and stalk area in sequential coronal sections along the rostrocaudal axis (n=9). * = p <0.05 ** = p<0.01, *** = p<0.001, One-way ANOVA. Error bars indicate standard error of the mean (SEM).

**Figure S1.**
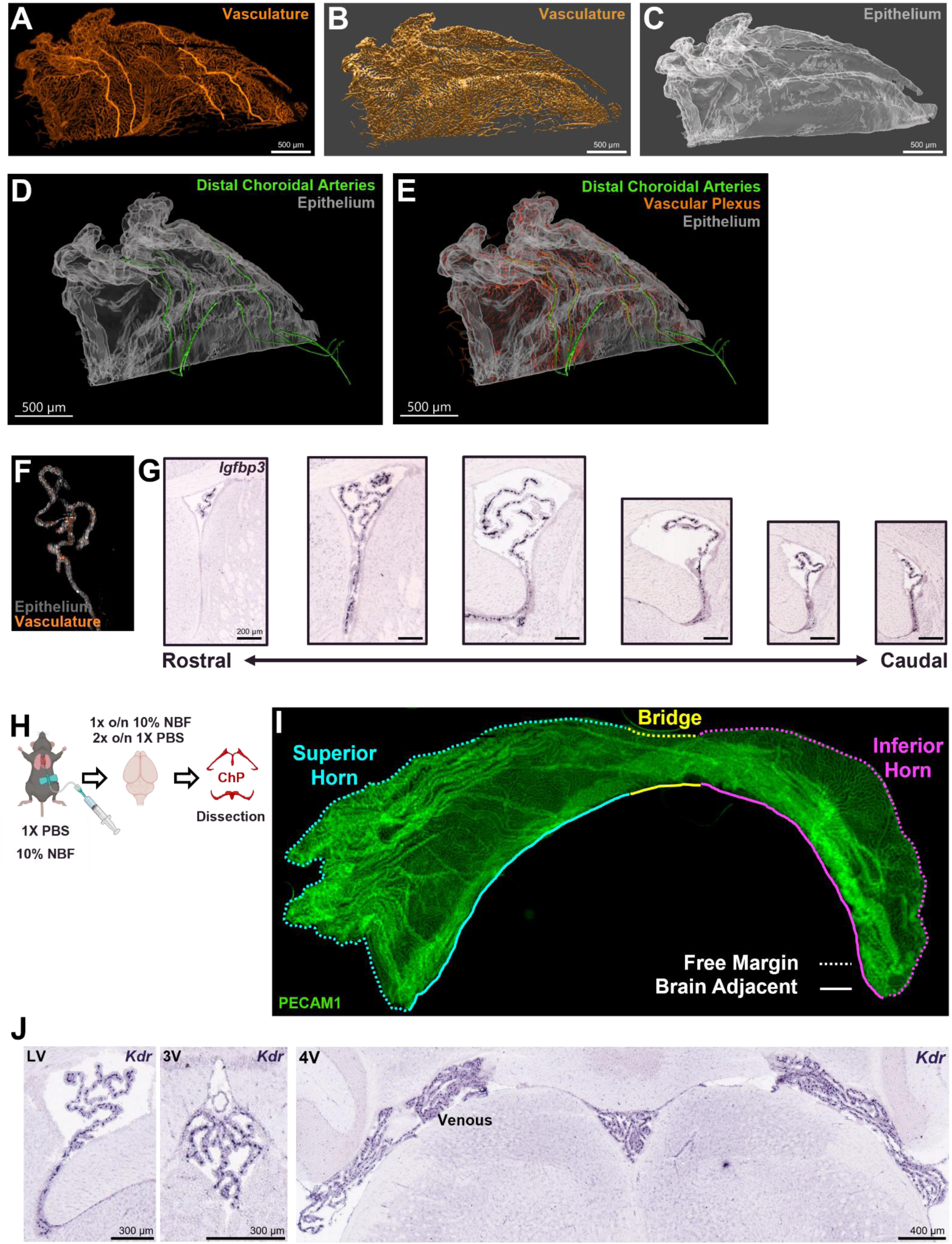
Choroid plexus structure in situ, ex vivo, and across ventricles. (A-C) Maximum projection micrograph of the choroid plexus that resides in the superior portion of the lateral ventricle (A) and Imaris 3D-renderings of superior horn vasculature (B) and epithelium (C). (D-E) 3D rendering of superior horn of the lateral ventricle choroid plexus. (F) Coronal optical section of the lateral ventricle of choroid plexus from *Tek-Cre::Rosa26^mT/mG^* transgenic mouse. Vasculature is represented in orange while non-endothelial tissue is grey. (G) Examples of rostral to caudal coronal sections at 100 µm used to quantify stalk and ventricular portions of lateral ventricle choroid plexus. Images show *in situ* hybridization of insulin-like growth factor-binding protein 3 (*Igfbp3*) expression in the lateral ventricle choroid plexus (Allen Brain Atlas). Scale bar is 200 µm. (H) Workflow of cardiac perfusion tissue processing protocol. (I) Lateral ventricle choroid plexus explant with labeled anatomical regions including superior horn (cyan), bridge (yellow), and inferior horn (magenta) have distinct morphologies with complex 3D morphology (free margin; dotted line) and planar brain adjacent regions (solid line). (J) Coronal sections of the lateral, third, and fourth ventricle of adult mouse brain.

### Ventricle and age associated gene expression profiles of choroid plexus endothelia

To investigate molecular underpinnings of endothelial architecture and function, we reanalyzed a published single-nucleus sequencing dataset from Dani et al. (GSE168704) to uncover the transcriptional basis of endothelial subtypes and identify rational targets for functional studies. Subsets of annotated endothelial nuclei from the lateral, third, and fourth ventricle choroid plexuses across embryonic day (E) E16.5, adult (4 months) and aged (20 months) mouse brains were analyzed in Seurat (**Fig. 2A**, see Methods). Nuclei could be partitioned into six different subclusters: Clusters 1, 4, and 5 mapped exclusively to embryonic mice while clusters 0, 2, and 3 were found in both adult and aged mice (**Fig. 2B, 2C**). Capillary markers were enriched in Cluster 0 (*Plpp1*, *Syne1*, *Jam3*) and Cluster 1 (*Pde4d*, *Airn*, *Tref1*), which mapped to embryonic and adult stages, respectively. Further, age-specific expression was found in Cluster 2 (*Vwf*, *Cfh*, *Pde3a)* and Cluster 3 (*Bmx*, *Nebl*, *Sema3g*), which marked venous and arterial subtypes from adult choroid plexus, while Cluster 4 (*Trpm3*, *Slc6a15*, *Glis3*) and Cluster 5 (*Top2a*, *Mki67*, *Cenpp*) were found only in embryonic samples (**Fig. 2D**). To identify biological processes encoded by these gene expression profiles, we performed Gene Ontology (GO) enrichment analysis across each endothelial subcluster (**Fig. 2E**).

**Figure 2.**
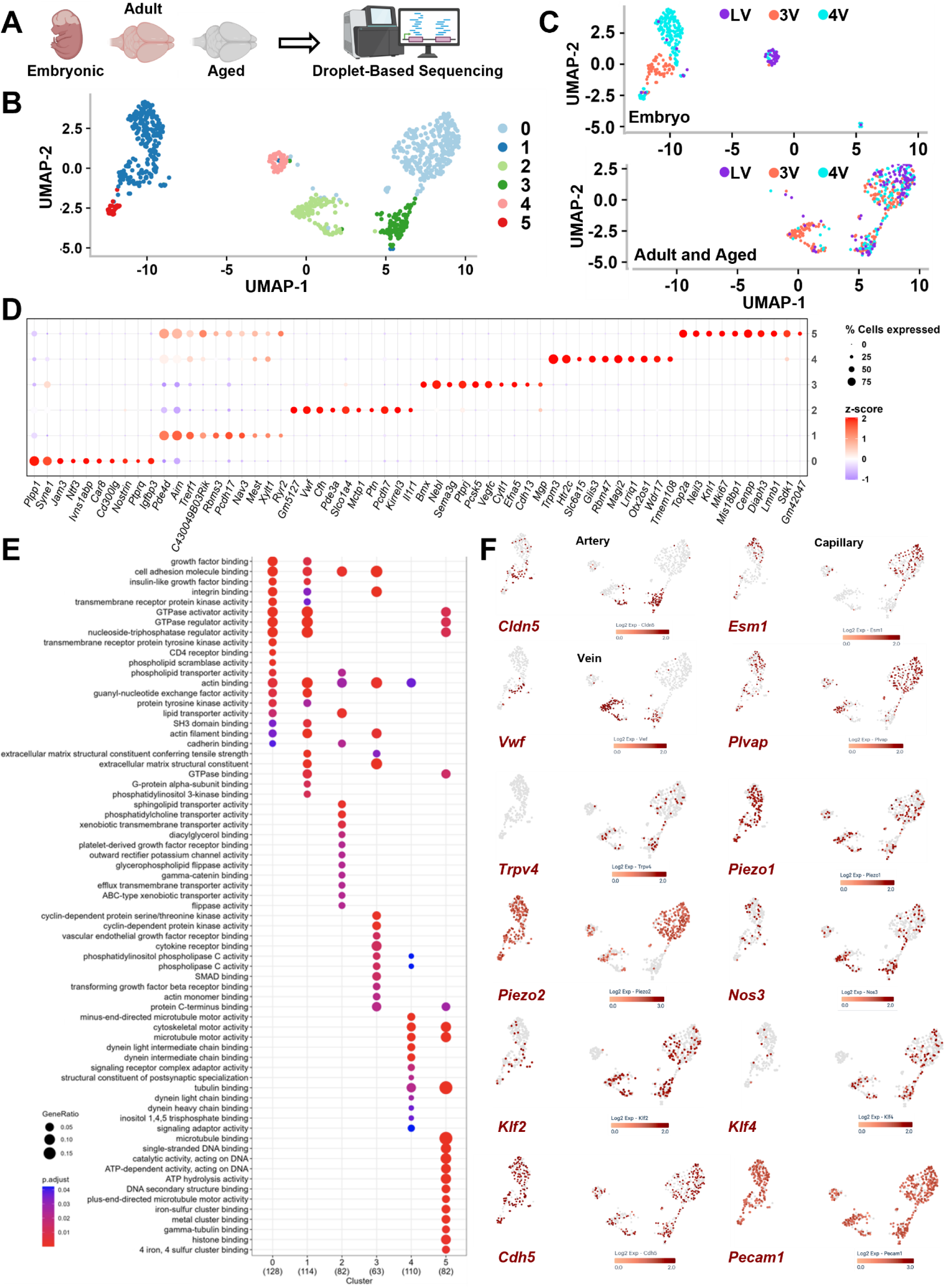
Transcriptional underpinnings of endothelial networks of the choroid plexus across brain regions and age. (A) Sequencing workflow. (B) UMAP plot shows endothelial subclusters. (C) Ventricle and age-specific clusters. Clusters 1, 4, and 5 are embryonic, while clusters 0, 2, and 3 are found in both adult and aged samples. (D) Dot plot with top-ranked genes in each cluster. (E) Dot plot displays the top enriched Gene Ontology (GO) terms (molecular functions) for each endothelial cluster identified by over-representation analysis using significantly expressed genes (p ≤ 0.05) within each cluster. Dot size represents the number of genes from a given cluster associated with each GO term, while dot color indicates the significance level of the enrichment (adjusted p-value). (F) UMAP plots of endothelial identity and mechanoresponsive gene expression.

**Figure S2.**
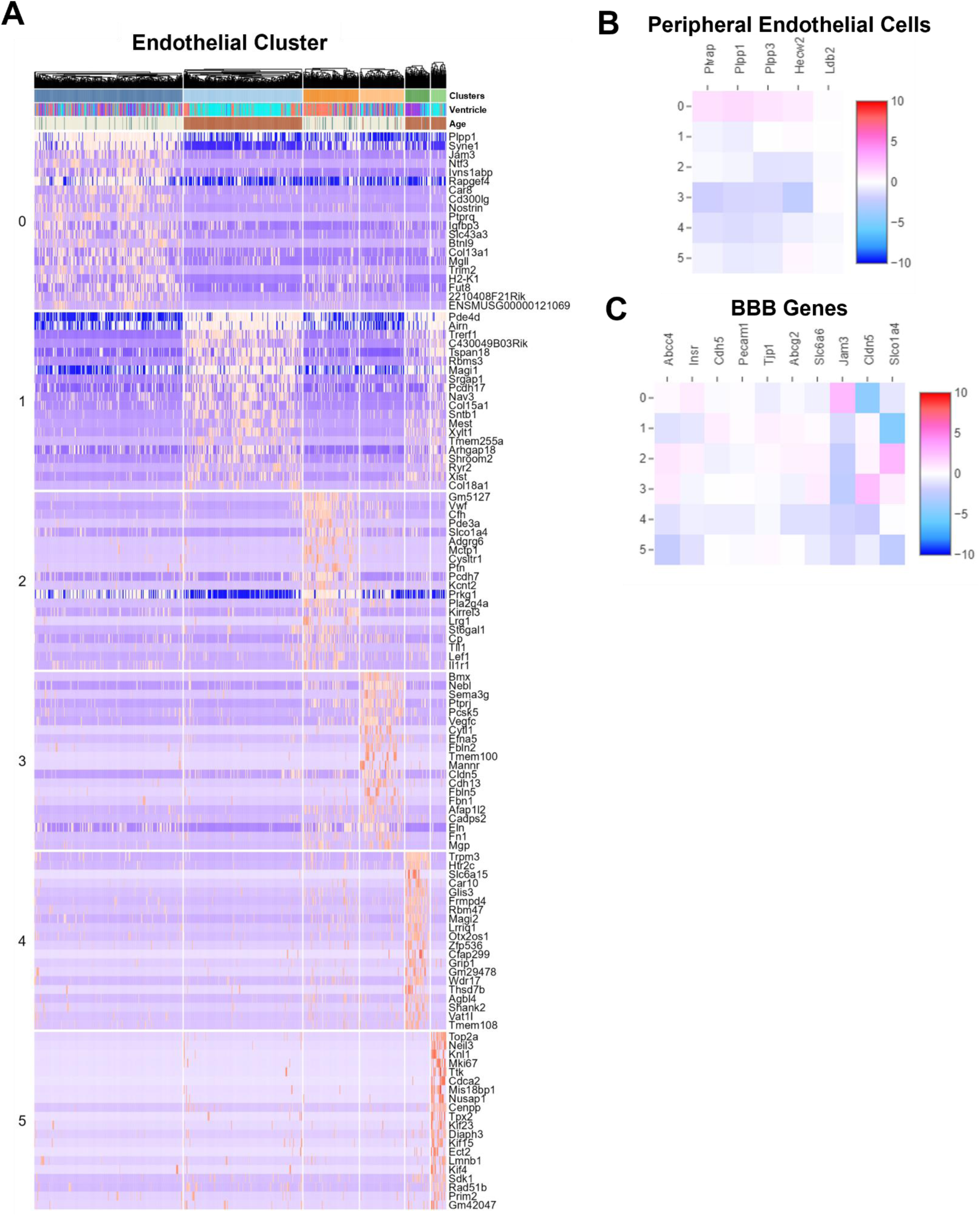
Endothelial gene expression of choroid plexus. (A) Heat map of differential gene expression across clusters, brain ventricles, and age. (B) Heatmap of peripheral endothelial cell gene expression across choroid plexus endothelial cell clusters. (C) Heatmap of canonical blood-brain barrier associated gene expression across endothelial cell clusters.

Capillary, arterial, and venous clusters strongly expressed cell adhesion molecules (e.g., *Itga2*, *Jam3*, *Vcam1*, *Kirrel3*), which confer junctional integrity and mechanical cohesion necessary for barrier tightness, vessel stability and controlled leukocyte traffic. Upregulated small-GTPase signaling (e.g., *Nrp1*, *Dlc1*, *Mcf21*, *Plcb1*) was found in embryonic and adult capillaries, which suggests the capillary endothelial network is primed for dynamic and signal-responsive regulation of cell adhesion, membrane composition, and cytoskeletal reorganization. This is consistent with reports that the choroid plexus endothelial cells form a vascular barrier with tunable permeability properties.^19^ The capacity for dynamic responses is further supported by expression of diverse actin binding proteins (e.g., *Syne1*, *Cobl*, *Rai14*, *Shank3*) that endow stable yet malleable actin architectures capable of nucleo-cytoskeletal coupling and finely tuned membrane organization. Next, embryonic endothelial cells clustered into two specialized subsets not found in other ages. One subset showed enriched kinesin, myosin, and dynein motor expression (e.g., *Kif6*, *Myo5b*, *Dnah7b*) while the other subset expressed mitotic spindle kinesins (e.g., *Kif11*, *Cenpe*), spindle organizers (e.g., *Tpx2*, *Ckap5*) and Rho/Rac pathway regulators (e.g., *Tiam1*, *Racgap1*), which identify dividing cell types. Embryonic capillary beds also upregulated phospholipid genes (e.g., *Plscr1*, *Plscr2*, *Plscr4*), which likely reflects a high capacity for tuning membrane lipid composition and phosphoinositide signaling for secretion, endocytosis, and stress response. In contrast, adult and aged capillary beds upregulated components for structural stabilization (e.g., *Col4a1*, *Lamc1*, *Nid2*) and growth-factor mediated regulation (e.g. *Igf1r*, *Igf2r*, *Gnas*) of adhesion strength. Adult and aged venous vessels appear optimized for lipid and xenobiotic transport (e.g., *Abcb1a*, *Abcc4*, *Slco1a4*, *Slc2a1*), ion channel excitability (e.g. *Kcnt2*, *Kcnb1*, *Unc13b*), and vascular permeability (e.g., *Ptprb*, *Ptprj*). Corresponding arterial endothelial cells show upregulated cell-matrix and intercellular adhesion markers (e.g., *Fbn1*, *Fbln1*, *Vcam1*, *Cdh13*) under control of growth factor signaling (e.g., *Vegfc*, *Tgfb2*, *Smad6*). Blood brain barrier genes (e.g., *Cldn5*, *Cdh5*) were primarily upregulated in subsets of adult and aged vessels, which were previously mapped to infiltrating endothelial cells that integrated with fenestrated capillary bed markers (e.g. *Plvap*, *Plpp1*, *Plpp3*) (**Fig. 1F**). Collectively, these data suggest the choroid plexus vascular bed is specialized, developmentally stratified, and mechanically robust with age-dependent remodeling capacity.

### Mechanosensitive calcium signaling in subsets of choroid plexus endothelial cells

To identify upstream regulators of endothelial dynamics we interrogated expression of key genes linked to hemodynamic mechanotransduction. We found embryonic and adult cell types expressed shear-responsive endothelial transcription factors (e.g., *Klf2*, *Klf4*) and mechanosensitive calcium channels (e.g. *Piezo1*, *Piezo2*, *Trpv4*) that are known to control endothelial cell alignment, vascular tone, and permeability (**Fig. 2F**). To visualize calcium activity in real time we crossed the endothelial driver line (Tek-Cre) with a transgenic calcium reporter mouse, enabling Cre-dependent expression of a genetically encoded calcium indicator, GCaMP, in endothelial cells. We aged endothelial-GCaMP (Tek-Cre::GCaMP6s) mice to adulthood, excised whole choroid plexus tissue and stabilized explants in a glass bottom dish using surgical glue (**Fig. S3A**). This preparation allows visualization of arterial, venous, and capillary endothelial cells across the entire choroid plexus while keeping local vascular connectivity and tissue microenvironment largely intact (**Fig.3A, 3B**). We identified entry points of distal choroidal arteries into the choroid plexus vascular beds and performed confocal imaging calcium spiking activity (see methods) (**Fig. 3A-C**). Earlier work showed that the infiltrating vessels are arteries that feed the capillary beds of the choroid plexus. Spontaneous calcium spiking events were most evident along these vessels and calcium activity transients were extracted from video recordings of regions along this vessel (see methods). Patterns of spiking activity appeared graded, as peak amplitude and frequency of events were higher in segments of the infiltrating vessel proximal to the brain as compared to distal segments extending towards the free margin of the tissue (**Fig. 3C, 3D**). Regions of high spiking activity showed rhythmic contractions consistent with high vasomotor activity (**Fig. 3E, 3F**). Plotting the distribution of spiking amplitude (dF/F) and luminal width over time showed that calcium spiking amplitude was highest in regions of vessel contractions (**Fig. 3G**). In regions of high activity, peak luminal contraction appeared to coincide with peak calcium amplitude, which underscores coupling of vascular tone and calcium activity in subsets of choroid plexus endothelial cells (**Fig. 3H**).

**Figure 3.**
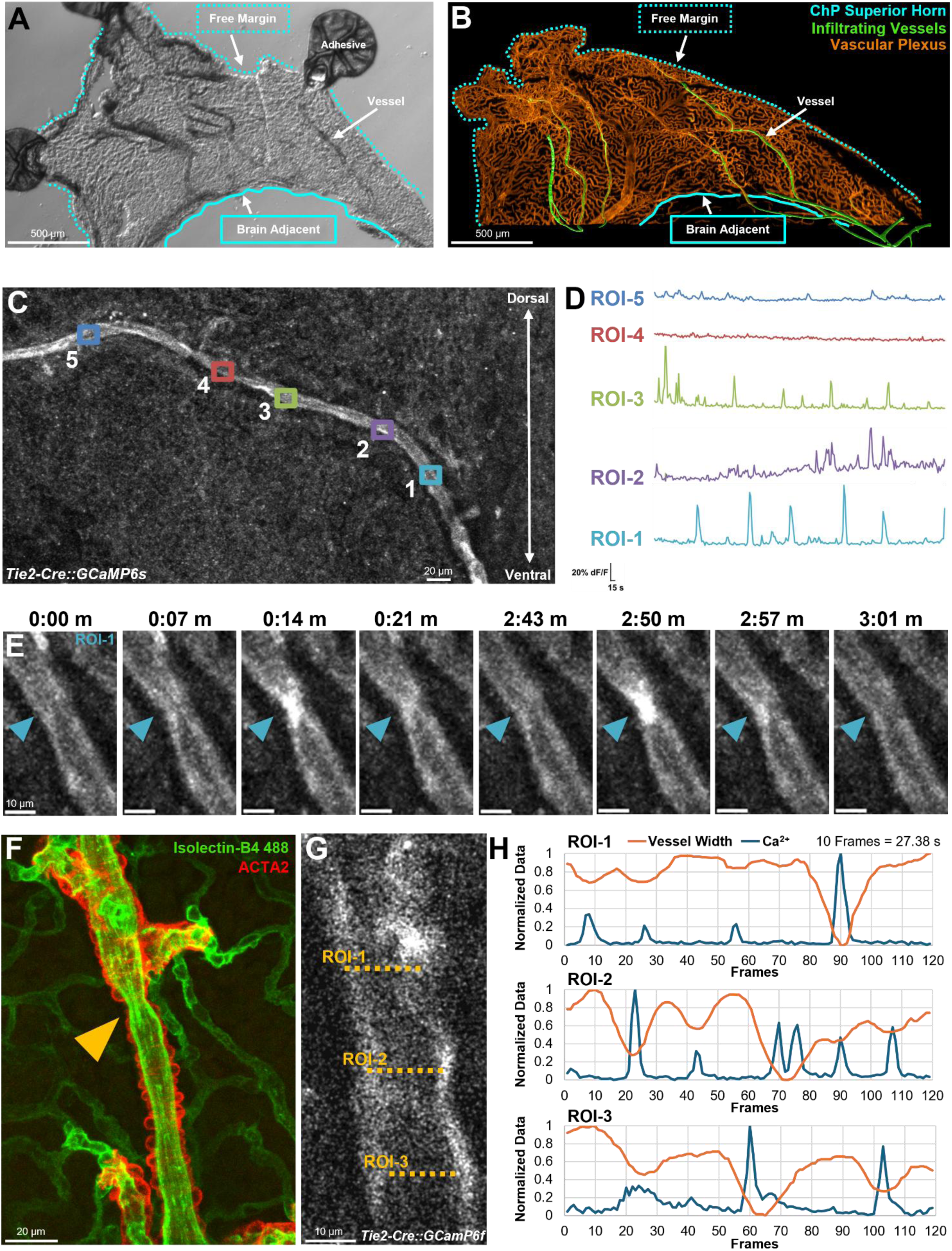
Explant preparation to measure spontaneous calcium activity in vessels. (A) Brightfield micrograph of a lateral ventricle choroid plexus explant secured with tissue-safe adhesive for stable calcium imaging. Dotted lines mark the free margin and solid lines brain adjacent tissue regions. (B) Imaris rendering of the vascular plexus highlight tissue infiltrating vessels in superior horn of lateral ventricle explant. (C) Maximum intensity projection of fluorescence over time of a 3D stack recorded from an endothelial-calcium reporter mouse. Colored squares identify regions of interest (ROIs) along a single arterial vessel. (D) Traces of calcium spiking events in ROIs along a single infiltrating vessel. (E) Snapshots of local calcium activity and rhythmic vessel contractions at ROI-1 that is closest to ventral connection points with brain. (F) Maximum intensity projection of a contracting artery (yellow arrow). Isolectin-B4 labels endothelia and smooth muscle α-2 actin (ACTA2) highlights ensheathing myofibroblasts. (G) A single imaging frame of GCaMP fluorescence in an endothelial vessel. (H) Traces map temporal changes in vessel width and calcium activity in ROIs identified in 3G.

**Figure S3.**
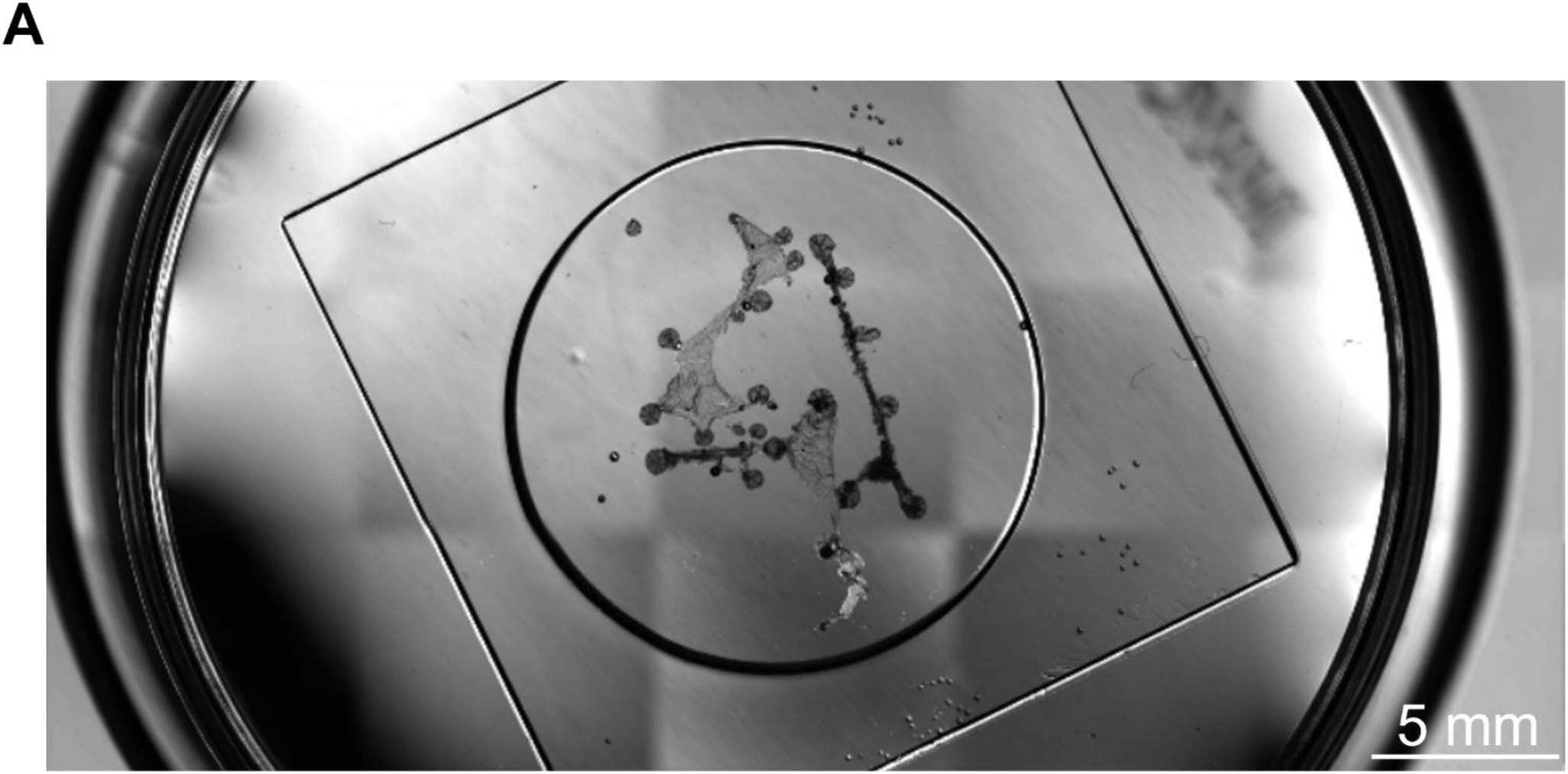
Explant preparation and stabilization of multiple choroid plexus explants. (A) Multiple choroid plexus explants in a glass-bottom dish.

### Piezo1-activated mechanotransduction triggers endothelial calcium dynamics and modulates cell adhesion

Brain endothelial cells use mechanotransduction mechanisms to build, maintain, and remodel blood-brain barrier properties and tune local responses to shear, pressure, and stretch forces. For example, shear sensing complexes at endothelial junctions (PECAM1, VE-cadherin, VEGFR2) convert flow into signaling that tightens endothelial junctions to lower permeability, while abnormal or pulsatile flow overactivate mechanosensitive channels (Piezo1, TRPV4) promoting junctional disassembly and BBB leakage.^32,34^ Mechanotransducing properties of the choroid plexus endothelia are largely unknown, are inferred from general endothelial studies and are vastly underappreciated compared to the BBB. To directly test mechanobiological responses of intact choroid plexus vascular beds, we employed a live imaging chamber that enabled perfusion of intact choroid plexus explants with mechanobiology-targeted pharmacology in a flowrate, temperature, oxygen, and humidity-controlled environment (See Methods) (**Fig. S4A**). Experiments testing perfusion rates using increasing concentrations of methylene blue in artificial cerebrospinal (aCSF) showed rapid exchange of bath contents (**Fig. S4A, S4B**). From mechanosensitive genes identified by single nucleus analysis (**Fig. 2F**), we prioritized investigation of the Piezo1 ion channel for its role in coupling calcium activity with mechanotransduction, availability of pharmacological tools, and limited prior investigation in vascular beds of the choroid plexus. To test Piezo1 calcium activity in choroid plexus endothelial cells, we employed a transgenic mouse line that constitutively expressed calcium indicator GCaMP8f in endothelial cells under the control of the VE-cadherin (Cdh5) promotor (**Fig. S4C**).

We dissected and stabilized lateral ventricle choroid plexus explants from embryonic and adult mice in a perfusion rig and live imaged calcium responses to a selective Piezo1 agonist, Yoda1 (**Fig. 4A, 4B**). First, tissue health was ascertained by looking for spontaneous calcium activity. Vessel segments with sustained high calcium levels along glue-stabilized edges, likely indicative of poor health, were excluded from analysis. Second, we looked for regions with spontaneous calcium spiking activity and found active regions in both embryonic and adult explants. In these ROIs, brief exposure to Yoda1 triggered robust calcium responses (**Fig. 4A-C**, See methods).^30^ Piezo1 activation evoked an increase in intracellular embryonic calcium levels (orange arrowheads), which appeared in a wave-like pattern that was sustained for several minutes (**Fig. 4A, Video S1**). In adult explants, Yoda1 evoked rhythmic calcium oscillations across and within endothelial segments (**Fig. 4B, Video S2**). To quantify changes in calcium responses before and after Yoda1 exposure we developed a custom algorithm that allowed pixel level classification of calcium transients (**Fig. 4C**). Averaging calcium intensity measurements from active endothelial tissue over time showed that both embryonic and adult explants evoked network-wide calcium responses to Yoda1 (**Fig. 4D-F, S4D**). Next, to test the consequence of Piezo1 activation on cell adhesion, we immunolocalized cell adhesion protein PECAM1 in pre- and post-flow conditions with and without Yoda1 (**Fig. 4H**). Little to no change was observed in the endothelial glycocalyx (Isolectin-IB4) and ETS-family transcription factor (ERG), which maintains endothelial identity, survival, and junctional gene expression (data not shown). In contrast, PECAM1 showed marked differences between explants that were immediately fixed and those subjected to flow conditions (**Fig. 4H**). Yoda1-treated explants showed PECAM1 fluorescence intensity and signal distribution more like freshly fixed tissue compared to untreated explants (**Fig. 4I**, * = p <0.05 ** = p<0.01, *** = p<0.001). This is consistent with reports that the agonist Yoda1 opens Piezo1 channels, effectively mimicking key aspects of shear forces that in choroid plexus endothelia stabilize cell adhesion.

**Figure 4.**
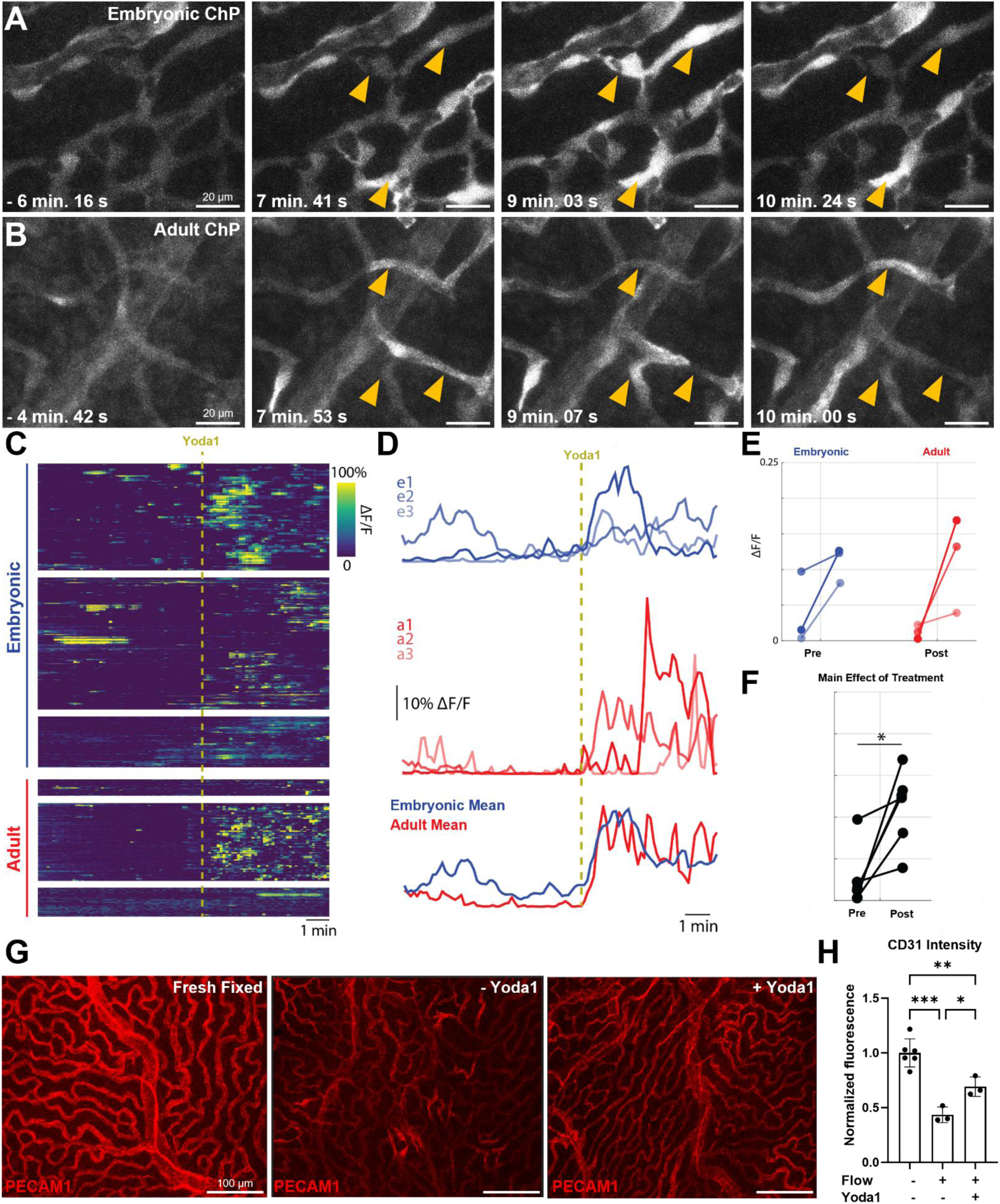
Piezo1 agonists trigger calcium dynamics and modulates cell adhesion. Snapshots of Yoda1 evoked calcium responses in GCaMP expressing embryonic (A) and adult (B) endothelial explants. (C) Heat map shows temporal changes in GCaMP fluorescence in embryonic and adult choroid plexus endothelial cells pre- and post Yoda1 stimulus. (D) Traces show individual average calcium spiking activity over time in embryonic and adult explants. (E-F) Quantitation of Yoda1 responses (n=3, embryonic and adult explants) reveals a main effect of treatment but not age. Two-factor ANOVA (* = p<0.05). (G) Representative images of PECAM1 distribution in vascular beds of freshly fixed, aCSF only, and Yoda1 treated explants. (H) Normalized mean fluorescence intensity of PECAM1 within vasculature across individual groups (fresh fixed, n=6, Yoda1 treated and untreated samples, n=3). * = p <0.05 ** = p<0.01, *** = p<0.001. One-way ANOVA with Tukey’s multiple comparisons test. Error bars indicate (SEM).

**Figure S4.**
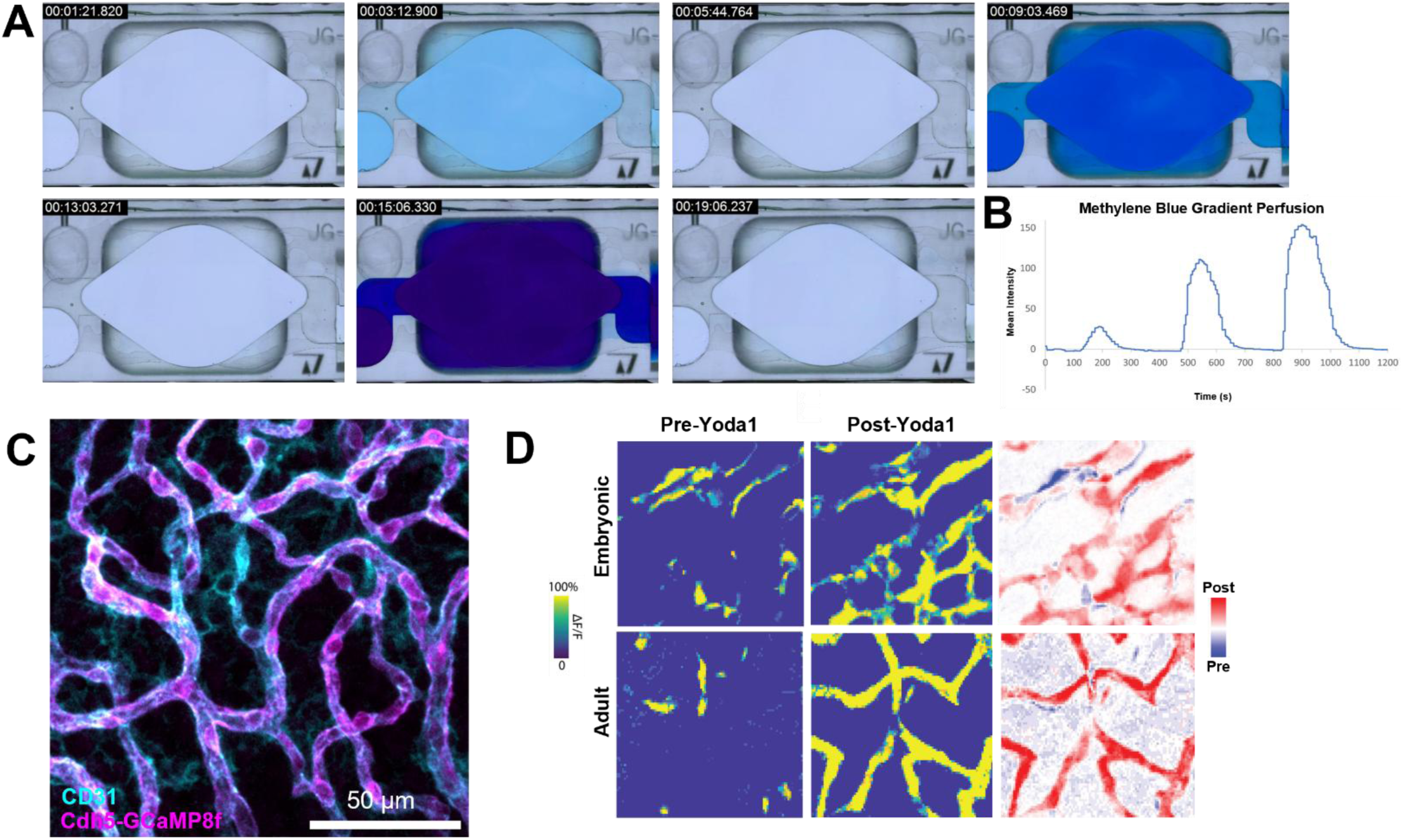
Choroid plexus flow chamber and calcium responses. (A) Snapshots show bright field images of progressively increasing concentrations of methylene blue and clearance under flow at 5 mL/min, (B) Quantitation of signal intensity over time, (C) Cdh5-GCaMP8f expression in fixed tissue explant. (D) Heatmaps show Yoda-1 evoked fluorescence intensity changes are localized to endothelial cells in embryonic and adult explants.

## DISCUSSION

The BBB has dominated cerebrovascular research for decades and investigation of other blood-CSF barriers including the meninges and circumventricular organs is substantially ahead of choroid plexus research. A growing number of studies highlight the specialized properties of choroid plexus endothelial cells that directly interface with blood content. They are the first line of defense against pathogens entering the choroid plexus and regulate traffic of peripheral immunity signals into the brain in models of autoimmune disease.^25^ Despite having natural apertures called fenestrae, these endothelial cells can form vascular barriers that integrate information coming from the gut and show dynamic permeability properties in WNT-signaling dependent manner.^19,26^ Therefore, choroid plexus barrier functions is likely multilayered and distributed between epithelial cell monolayers and the underlying vascular plexus. To fill this gap, we present a multimodal tool kit and provide a framework for systematic interrogation of endothelial gene functions that regulate structural and functional properties. First, we reasoned that mapping structural features of the vascular bed and connections with rest of the cerebrovasculature would reveal how choroid plexus vessels are integrated with global brain blood flow and identify routes for signal, solute, and immune cell exchange between the blood and CSF. Earlier light sheet imaging studies employed antibody-based labeling strategies focused on parenchymal vasculature, and investigation of fine-scale organization as seen in the choroid plexus was limited to large arteries and venous outflows.^43,45^ Here, we employed a transgenic double fluorescence labeling strategy that drives fluorescent molecules (EGFP, tdTomato) to membranes of endothelial and non-endothelial cell types, which enables high-resolution imaging of local vascular connectivity and integration of the choroid plexuses with the rest of the cerebrovasculature. Consistent with earlier vascular mapping studies that identified molecularly specialized vessels (*Cldn5*+) in explants, we identified *in vivo* orientation of distal choroidal arteries that enter the superior horn of the lateral ventricle choroid plexus. Our dataset will enable mapping of similar influx and efflux pathways across choroid plexuses in each ventricle at high-resolution in the adult mouse brain. As with other approaches, we found tissue clearing and image quality was reduced in deep brain structures. Emerging optical strategies including adaptive optics-assisted light-sheet microscopy, two-photon light-sheet imaging, and expansion microscopy combined with volumetric imaging may improve light penetration and provide deeper reconstruction of fine vascular structures. Integration with spatial transcriptomics profiling and deep learning-based segmentation will offer improved opportunities for future integrative studies of vascular networks.

Earlier bulk and single-cell transcriptomic studies have reported ventricle- and age-specific enrichment of epithelial and fibroblast gene expression, which is thought to confer regionalized functional specialization and local signaling with adjacent brain regions. Here, we find that endothelial gene expression is most divergent between embryonic and adult stages. Transcriptional pathway analysis suggests that choroid plexus vascular beds transition from lipid- and motor-rich, highly plastic embryonic endothelium to a structurally reinforced, segment-specialized network in adults. However, G-protein coupled receptors and growth-factor receptors were found in embryonic, adult, and aged choroid plexus endothelia, which suggests the vascular bed remains responsive to diverse stimuli across lifespan. We also found vessel and age specific expression of several receptor pathways and intracellular machinery known to drive vasoactive responses to increased blood flow, pressure, pain, and inflammation. Coupled with our live imaging workflow, these transcriptional insights will enable testing of choroid plexus vascular-specific responses that may regulate fenestrae dynamics and cell-cell interactions with resident cell types that distinguish properties from other brain barriers.

Calcium imaging studies have been performed on primary brain endothelial cultures, retinal microvascular beds, and *in vivo* to interrogate barrier properties, vasodilation, sprouting angiogenesis, inflammation, and neurovascular coupling. However, no earlier studies have performed any investigation of calcium linked endothelial properties in the choroid plexus endothelium. In our novel imaging preparation that retains vascular connectivity of resident endothelial cells, we found spontaneous and regionalized calcium oscillations in subsets of arterial endothelial cells. These events were found along transition zones between the brain and choroid plexus vascular beds, which likely reflect spontaneous mechanisms to pump blood into the choroid plexus vascular network. In addition to spontaneous activity, we found strong evoked responses in embryonic and adult endothelial cells, with distinct intra- and intercellular activity patterns in response to Yoda1, a Piezo1 agonist. In BBB, Piezo1 regulates intravascular force sensing during functional hyperemia, barrier properties in ischemia, flow-dependent vascular development, and interactions with pathological ligands in neurodegeneration.^36,38,41^ Here, we reasoned to investigate effects on cell adhesion, as classic work has shown that elevation of intracellular calcium activity is tied to junctional adhesion.^49^ We found that Piezo1 ion channels can stabilize PECAM1 expression in *ex vivo* conditions, which contrasts with Yoda1 perfusion *in vivo*, which led to disordered PECAM1 patterning in retinal vasculature.^27^ One explanation is that Yoda1 injections over activate shear sensing pathways leading to disruption of cell adhesion. In the same study, defects were only observed in venous vessels and not arteries of the retinal vasculature indicative of subtype or tissue-specific responses. A caveat of the explant preparation is the lack of luminal blood flow known to be important for endothelial health and barrier function. This was reflected in our explant imaging studies, where glycocalyx and ERG expression remain largely unchanged, but PECAM1 distribution at cell-cell junctions appeared to be irregular. PECAM1 expression was partially rescued by acute stimulation (1 min) by Yoda1, which functions as a pharmacological mimetic of shear stress. Partial rescue could be explained by the presence of multiple cell adhesion mechanisms, where loss of one endothelial adhesion system is buffered by others (e.g., VE-cadherin).^51^ While innovations in choroid plexus intravital imaging will offer rich *in vivo* insights, these studies will be limited by 2-photon access only to exposed regions around the free margin of the lateral ventricle choroid plexus. In contrast, explant approaches allow imaging access to vessel networks across the entire choroid plexus from any ventricle and developmental stage without the need for invasive surgical approaches. Our framework can be readily adapted to include investigation of other blood-CSF barriers including the meninges and CVOs, providing broader accessibility and ease of use at relatively low cost.^39^ Overall, our work shows previously unappreciated structural and functional dynamics of choroid plexus vascular beds and motivates further investigation of calcium signaling pathways tied to barrier function and vasodilation in this blood-CSF barrier. This work synergizes with growing interest in probing cell-cell interactions that regulate local endothelial, mural, neural, glial, and immune microenvironments in the brain.^46–48,50^

## METHODS

### Experimental Models

All animal care and experimental procedures were approved by the Institutional Animal Care and Use Committees of Vanderbilt University and Vanderbilt University Medical Center. Mouse lines used include mT/mG (Jax#: 007676, RRID:IMSR_JAX:007676), Tek-Cre (Jax#:008863, RRID:IMSR_JAX:008863), Ai96(RCL-GCaMP6s) (Jax#: 024106), Cdh5-GCaMP8f (Jax#: 033342, RRID:IMSR_JAX:033342). All animals were housed under 12hr/12hr day night cycle with access to standard chow and water ad libitum.

### Brain tissue clearing

Mouse brain tissue was prepared according to the LifeCanvas SHIELD protocol (Stabilization under Harsh conditions via Intramolecular Epoxide Linkages to prevent Degradation) for the immobilization of proteins. Active clearing was performed by delipidation using SmartClear+ (LifeCanvas) which uses stochastic electrotransport to remove electromobile molecules, such as phospholipids, from the sample, following the SmartClear+ User preset for clearing (limit = 90 V, current = 1500 mA, temp. = 42 °C)..

### Light sheet imaging

To generate fluorescently labeled endothelial cells in mice we crossed Tek-Cre males with mT/mG females. Three-dimensional imaging was performed on a SmartSPIM single-plane illumination light sheet microscope (LifeCanvas Technologies) equipped with a 3.6x objective lens with dipping cap (NA 0.2, uniform axial resolution approximately 4 µm) and an sCMOS camera with a rolling shutter.^29,31^ Samples to be illuminated with two excitation wavelengths: 488 and 561 nm. Emissions detected with 525/50, and 600/52 filters. Following acquisition, images are transferred to a Dell Precision 7920 Tower with (2) Intel Xeon Gold 6248R CPU at 3.0 GHz and 512 GB of RAM running Windows 11 Pro for Workstations for stitching and processing. Raw composite TIFF images geneMouseed by the camera are automatically de-stripped and stitched using a modified implementation of Terastitcher (Bria et al., 2019) from LifeCanvas Technologies. Processed TIFF stacks will be converted to Imaris (Bitplane, Oxford Instruments) files using Imaris File Converter 9.9.0. Volumetric renderings, keyframe animations and digital segmentation to be performed in Imaris v10.2. Image resolution of 2umx2umx1.8um provide near isotropic images.

### Imaris and 3D rendering

Three-dimensional image stacks containing the lateral ventricle and choroid plexus (CP) were processed using Imaris version 10.2 (Oxford Instruments). The lateral ventricular region containing the CP was first isolated by manual cropping in three dimensions to reduce background signal and restrict subsequent segmentation to the ventricular compartment. A manual mask was generated to isolate the CP signal prior to surface reconstruction. CP surfaces were generated using the Imaris Surface module. Surface smoothing was enabled with a surface grain size of 5.40 µm to reduce high-frequency noise. Background elimination was not applied to preserve low-intensity CP structures. Surface detection was performed using a manually defined absolute intensity threshold (threshold value = 900). Following surface generation, objects smaller than 250 voxels were excluded to remove residual noise and sub-resolution structures. The resulting CP surface object was exported from Imaris as a Virtual Reality Modeling Language (WRL) mesh file for downstream visualization and geometric analysis. The exported surface was imported into MeshLab (version 2025.07) for rendering and surface-based visualization.

Large vessels were segmented independently using the Imaris Surface module. Surface smoothing was enabled with a surface grain size of 3.60 µm to preserve vessel continuity while reducing segmentation noise. Machine learning pixel classification within Imaris was trained using a single vascular channel. Following segmentation, surface objects smaller than 500 voxels were excluded to remove small or discontinuous vascular fragments. To restrict analysis to vasculature directly associated with the CP, a spatial filtering step was applied using the Imaris distance-to-surface metric. Only vessel surfaces with a shortest distance of less than 1 µm to the CP surface were retained.

### Ventricle to Stalk Area quantitation

Images of consecutive coronal sections at 100 µm distances were collected from Allen Brain Atlas murine brain atlas at postnatal day (P) P56 from 9 different gene sets (Efna5, Igf2, Kdr, Cldn5, Flt1, Lyve1, Vwf, Pptrj, Igfbp3). Choroid plexus tissue in the lateral ventricle was designated as ventricle area while stalk region was defined as the portion of the tissue in the choroidal fissure. Statistical analysis of data included Brown-Forsythe ANOVA tests with Tukey’s multiple comparisons test.

### Single-nuclei RNA-seq Data Processing and Analysis

Single-nucleus RNA-seq (snRNA-seq) data from the choroid plexus were obtained from the Gene Expression Omnibus (GEO; accession number GSE168704, Dani et al. 2021). The dataset includes nuclei isolated from the lateral, third, and fourth ventricles of embryonic (E16.5), adult (4 months) and aged (20 months) mouse brains (BL6/J), collected in biological triplicates. Raw FASTQ files were processed using an Nextflow workflow for snRNA-seq preprocessing documented in https://github.com/porchard/snRNAseq-NextFlow. Alignment and gene quantification were performed using STARsolo in GeneFull_ExonOverIntron mode, which prioritizes exon alignments. The mm39 reference genome was used for alignment and Gencode M31 primary assembly annotations were used for gene quantification.

Further downstream preprocessing was performed in R (v4.1.1) using the Seurat R package (v 4.3.0). The cells were filtered on number of transcripts on “per sample” basis and considering dropkick quality control metrics.^33^ Additional filters were applied to retain high-quality cells (at least 500 genes per cell and less than 10% mitochondrial gene content). Raw counts for each cell were scaled by the total UMI count, multiplied by a scale factor of 10,000, and log-transformed to generate normalized expression values. To identify genes with high biological variability across cells, we applied the variance-stabilizing transformation (VST) method. The number of variable genes selected was set to the median number of detected features per cell in the dataset. After variable feature selection, expression values were scaled as z-scores. Principal component analysis (PCA) was performed using on variable genes out to 50 principal components. An elbow plot was used to determine the number of informative principal components (PCs), and the first 15 PCs were selected for downstream analysis. Cells were then embedded in a shared nearest neighbor (SNN) graph using and clustered at a resolution of 0.2 units. Finally, low-dimensional visualization was generated by applying UMAP using the same selected PCs. Ambient RNA contamination was estimated and corrected using SoupX in two steps: automated contamination estimation followed by manual adjustment of the contamination fraction per sample.^35^ Doublets were identified and removed using scDblFinder, with doublet scores computed from raw counts using sample IDs and initial clustering as inputs with expected doublet rate at 0.3. Harmony (Korsunsky et al., 2019) integration was applied accounting for sample ID and age variables.^37,40^ Cell type annotations were performed assessing overlap of cell type canonical gene marker expression in clusters.

Cells annotated as Endothelial (n = 1,616) were subset followed by normalization, scaling and clustering in similar manner as described above for full dataset. Two clusters expressing noisy epithelial signatures were removed as ambient contamination. Clustering was performed without batch correction using the first 20 PCs. A shared nearest neighbor (SNN) graph was constructed and clustering was performed at a resolution of 0.2. UMAP embeddings were generated using the same 20 PCs. Selected genes were visualized for age and ventricle variation via ComplexHeatmap (v2.15.4).^42^

Endothelial cluster-specific marker genes were identified by Seurat’s FindAllMarkers() using Wilcoxon Rank Sum test with parameters “only.pos = TRUE” and a minimum detection threshold of 25%. Genes with adjusted p-values < 0.05 were retained and grouped by cluster for downstream gene ontology enrichment. Gene symbols were converted to Ensembl and Entrez identifiers using clusterProfiler::bitr() (v4.10.1. Wu et al., 2021) with annotation databases (org.Mm.eg.db), and unmapped genes were excluded. Cluster-wise Entrez gene lists were subjected to Gene Ontology (Molecular function) enrichment analysis using clusterProfiler with over-representation testing and a p-value cutoff of 0.05 and semantic similarity among enriched GO terms was used to reduce redundancy.^44^

### Choroid Plexus Explant Preparation

Whole choroid plexus from the lateral ventricle was isolated using #5 forceps (Dumont, Cat. #: 11251-10) and a scalpel (Ambler Surgical, Cat. #: 961501) in cold 1X artificial cerebrospinal fluid (aCSF; 119 mM NaCl, 2.5 mM KCl, 26 mM NaHCO3, 1 mM NaH2PO4, 11 mM Glucose, 2 mM MgCl2, 2.8 mM CaCl2) in a SYLGARD (Dow, Cat. #: 04019862) coated dissection dish. To process the brain for lateral ventricle choroid plexus isolation, the hindbrain and cerebellum were first separated from the mid- and forebrain via a ventromedial scalpel slice. Hemispheres were then separated via a midline cut along the corpus callosum before the removal of approximately one-third of each hemisphere’s rostral end. Finally, the folded structure of the hippocampus was rolled out using two forceps and the lateral ventricle choroid plexus was gently separated from the fornix. Whole lateral ventricle choroid plexus explants were then transferred onto round, glass coverslips (15 mm, CELLTREAT Scientific Products, Cat. #: 229172) which a small dome of cold, 1X aCSF. The lateral ventricle choroid plexus explant was oriented on the round, glass coverslip such that tissue folding was reduced, and the explant remained flat. The explant was then secured to the round, glass coverslip with Vetbond tissue adhesive (3M, Cat. #: 1469SB). Anchor points of adhesive were dotted around the periphery of the explant such that the tissue adhered to the coverslip while being pulled taut. Successive rounds of additional adhesive were added to the explant periphery until the entire tissue was surrounded with adhesive. This ensured stabilization for sensitive live imaging. Secured samples were then moved to the imaging chamber.

### Ex vivo Calcium Imaging Experiments

Scanning confocal microscopy was used to record calcium activity in which choroid plexus endothelial cells express GCaMP6s (Tek-Cre::GCaMP6s) and GCaMP8f (Cdh5-GCaMP8f mice) using a Leica Mica Microhub scanning confocal microscope). Imaging was performed with a 10x, 0.32 NA (PL FLUOTAR, Dry) and 20x, 0.75 NA (HC PL ADO, CS2 Dry). LED power was optimized via Leica’s patented OneTouch technology. Three-dimensional (3D) recordings were performed in conjunction with the above settings with a consistent axial (z) step size (1.00 µm). The secured explant/coverslip complex was placed into the custom perfusion chamber with 1X aCSF (37°C) flowing. Subsequently, increasing concentrations of Yoda1 in 1X aCSF were introduced for 1-minute per concentration, with 15-minute 1X aCSF washouts in between treatments. To measure bulk tissue fluorescence, a maximum volume projection along the z dimension was performed to flatten each 3D volume into a 2D image, resulting in a 2D video across time. Vessel width was calculated after using gaussian blur functions on GCaMP6s fluorescence (ImageJ) followed by binarization and measurement of width reported as normalized values.

### Post-Live Imaging Immunofluorescence

To visualize protein expression and tissue structure changes post-ex vivo imaging, the secured explant/coverslip complex was removed from the perfusion chamber and placed in a spot dish to fix in 1.0 mL Bouin’s fixative (Electron Microscopy Sciences, Cat. #: 15990) for 10 minutes at room temperature. The explant was then subject to a rinse(R)-wash(W) series in 1X PBS (RRRW, 10-minute wash; RRW, 10-minute wash; RRW, 10-minute wash). A rinse is defined as the addition and immediate removal of a wash solution, such as 1X PBS. The fixed explant was gently freed from the adhesive securing it to the round, glass coverslip via a scalpel and forceps after the first three rinses. 3D image stacks were collected on Leica Mica Microhub scanning confocal microscope. Endothelial masks were generated by tracing out ROIs on binarized images. Immunofluorescence analysis included measurement of mean fluorescence intensity within entire vascular ROI. Statistical analysis of data included one-way ANOVA tests with Tukey’s multiple comparisons test.

### Statistical Analysis

All descriptions of statistical significance, statistical tests used, and exact values and representations of n can be found in the figure legends. Statistical analysis was performed on PRISM.

### Calcium transient measurements and activity changes in Piezo1 experiments

Movies were first registered using the NoRMCorre plugin in MATLAB. In FIJI, image resolution was downsampled 10x and a ΔF/Fb was calculated, where Fb was calculated using 3D median filter (x = 0, y = 0, z = 25). We wrote a custom-script in MATLAB that analyzed the ΔF/Fb of each pixel. We first identified all active endothelial tissue by thresholding pixel data based on two independent activity scores (variance of the data and signal to noise). Only pixels with high variance and high signal to noise, both measures that calcium transients occurred, were deemed active. Next, we used a Pearson correlation and hierarchical clustering to identify pixels with similar activity. The mean of endothelial tissue was then calculated before and after application of Yoda1. Means were compared using a two-factor ANOVA (factor 1: pre/post, factor 2: embryonic/adult).

## REFERENCES

1. Lun, M.P., Monuki, E.S., and Lehtinen, M.K. (2015). Development and functions of the choroid plexus-cerebrospinal fluid system. Nat. Rev. Neurosci. 16, 445–457. 10.1038/nrn3921.

2. Ya’el, C., Hochstetler, A., and Lehtinen, M.K. (2025). Choroid Plexus Pathophysiology. Annu. Rev. Pathol. 20, 193–220. 10.1146/annurev-pathmechdis-051222-114051.

3. MacAulay, N. (2021). Molecular mechanisms of brain water transport. Nat. Rev. Neurosci. 22, 326–344. 10.1038/s41583-021-00454-8.

4. Fame, R.M., and Lehtinen, M.K. (2020). Emergence and Developmental Roles of the Cerebrospinal Fluid System. Dev. Cell 52, 261–275. 10.1016/J.DEVCEL.2020.01.027.

5. Cui, J., Xu, H., and Lehtinen, M.K. (2021). Macrophages on the margin: choroid plexus immune responses. Trends Neurosci. 44, 864–875. 10.1016/J.TINS.2021.07.002.

6. Xu, H., Hehnly, C., and Lehtinen, M.K. (2025). The choroid plexus: a command center for brain–body communication during inflammation. Curr. Opin. Immunol. 93. 10.1016/j.coi.2025.102540.

7. Bitanihirwe, B.K.Y., Lizano, P., and Woo, T.U.W. (2022). Deconstructing the functional neuroanatomy of the choroid plexus: an ontogenetic perspective for studying neurodevelopmental and neuropsychiatric disorders. Molecular Psychiatry 2022 27:9 27, 3573–3582. 10.1038/s41380-022-01623-6.

8. Vara-Pérez, M., and Movahedi, K. (2025). Border-associated macrophages as gatekeepers of brain homeostasis and immunity. Immunity 58, 1085–1100. 10.1016/j.immuni.2025.04.005.

9. Reynolds, J.A., and Putterman, C. (2025). The choroid plexus in inflammatory and degenerative diseases of the central nervous system. Curr. Opin. Immunol. 95, 102588. 10.1016/j.coi.2025.102588.

10. Dani, N., Herbst, R.H., McCabe, C., Green, G.S., Kaiser, K., Head, J.P., Cui, J., Shipley, F.B., Jang, A., Dionne, D., et al. (2021). A cellular and spatial map of the choroid plexus across brain ventricles and ages. Cell 184, 3056–3074.e21. 10.1016/J.CELL.2021.04.003,.

11. Pellegrini, L., Bonfio, C., Chadwick, J., Begum, F., Skehel, M., and Lancaster, M.A. (2020). Human CNS barrier-forming organoids with cerebrospinal fluid production. Science 369. 10.1126/SCIENCE.AAZ5626.

12. Yang, A.C., Kern, F., Losada, P.M., Agam, M.R., Maat, C.A., Schmartz, G.P., Fehlmann, T., Stein, J.A., Schaum, N., Lee, D.P., et al. (2021). Dysregulation of brain and choroid plexus cell types in severe COVID-19. Nature 595, 565. 10.1038/S41586-021-03710-0.

13. Lun, M.P., Johnson, M.B., Broadbelt, K.G., Watanabe, M., Kang, Y.-J., Chau, K.F., Springel, M.W., Malesz, A., Sousa, A.M.M., Pletikos, M., et al. (2015). Spatially heterogeneous choroid plexus transcriptomes encode positional identity and contribute to regional CSF production. J. Neurosci. 35, 4903–4916. 10.1523/JNEUROSCI.3081-14.2015.

14. Lochhead, J.J., Yang, J., Ronaldson, P.T., and Davis, T.P. (2020). Structure, Function, and Regulation of the Blood-Brain Barrier Tight Junction in Central Nervous System Disorders. Front. Physiol. 11. 10.3389/fphys.2020.00914.

15. Sweeney, M.D., Kisler, K., Montagne, A., Toga, A.W., and Zlokovic, B. V. (2018). The role of brain vasculature in neurodegenerative disorders. Nat. Neurosci. 21, 1318–1331. 10.1038/s41593-018-0234-x.

16. Lacoste, B., Prat, A., Freitas-Andrade, M., and Gu, C. (2025). The Blood–Brain Barrier: Composition, Properties, and Roles in Brain Health. Cold Spring Harbor Perspectives in Biology 17. 10.1101/cshperspect.a041422.

17. Engelhardt, B., and Sorokin, L. (2009). The blood-brain and the blood-cerebrospinal fluid barriers: Function and dysfunction. Semin. Immunopathol. 31, 497–511. 10.1007/s00281-009-0177-0.

18. Schaeffer, S., and Iadecola, C. (2021). Revisiting the neurovascular unit. Nat. Neurosci. 24, 1198–1209. 10.1038/s41593-021-00904-7.

19. Carloni, S., Bertocchi, A., Mancinelli, S., Bellini, M., Erreni, M., Borreca, A., Braga, D., Giugliano, S., Mozzarelli, A.M., Manganaro, D., et al. (2021). Identification of a choroid plexus vascular barrier closing during intestinal inflammation. Science (1979). 374, 439–448. 10.1126/science.abc6108.

20. Miyata, S. (2015). New aspects in fenestrated capillary and tissue dynamics in the sensory circumventricular organs of adult brains. Front. Neurosci. 9, 390. 10.3389/fnins.2015.00390.

21. Muzumdar, M.D., Tasic, B., Miyamichi, K., Li, N., and Luo, L. (2007). A global double-fluorescent Cre reporter mouse. Genesis 45, 593–605. 10.1002/dvg.20335.

22. Stan, R. V., Tse, D., Deharvengt, S.J., Smits, N.C., Xu, Y., Luciano, M.R., McGarry, C.L., Buitendijk, M., Nemani, K. V., Elgueta, R., et al. (2012). The diaphragms of fenestrated endothelia – gatekeepers of vascular permeability and blood composition. Dev. Cell 23, 1203. 10.1016/j.devcel.2012.11.003.

23. Zagorska-Swiezy, K., Litwin, J.A., Gorczyca, J., Pitynski, K., and Miodonski, A.J. (2008). The microvascular architecture of the choroid plexus in fetal human brain lateral ventricle: a scanning electron microscopy study of corrosion casts. J. Anat. 213, 259. 10.1111/j.1469-7580.2008.00941.x.

24. Chang, T.H., Hsieh, F.L., Gu, X., Smallwood, P.M., Kavran, J.M., Gabelli, S.B., and Nathans, J. (2023). Structural insights into plasmalemma vesicle-associated protein (PLVAP): Implications for vascular endothelial diaphragms and fenestrae. Proc. Natl. Acad. Sci. U. S. A. 120, e2221103120. 10.1073/pnas.2221103120.

25. Kallal, N., Hugues, S., and Garnier, L. (2024). Regulation of autoimmune-mediated neuroinflammation by endothelial cells. Eur. J. Immunol. 54, 2350482. 10.1002/eji.202350482.

26. Wang, Y., Sabbagh, M.F., Gu, X., Rattner, A., Williams, J., and Nathans, J. (2019). Beta-catenin signaling regulates barrier-specific gene expression in circumventricular organ and ocular vasculatures. Elife 8. 10.7554/eLife.43257.

27. Chuntharpursat-Bon, E., Povstyan, O. V., Ludlow, M.J., Carrier, D.J., Debant, M., Shi, J., Gaunt, H.J., Bauer, C.C., Curd, A., Simon Futers, T., et al. (2023). PIEZO1 and PECAM1 interact at cell-cell junctions and partner in endothelial force sensing. Communications Biology 2023 6:1 6, 358-. 10.1038/s42003-023-04706-4.

28. Park, Y.G., Sohn, C.H., Chen, R., McCue, M., Yun, D.H., Drummond, G.T., Ku, T., Evans, N.B., Oak, H.C., Trieu, W., et al. (2018). Protection of tissue physicochemical properties using polyfunctional crosslinkers. Nature Biotechnology 2018 37:1 37, 73–83. 10.1038/nbt.4281.

29. Dean, K.M., Roudot, P., Welf, E.S., Danuser, G., and Fiolka, R. (2015). Deconvolution-free Subcellular Imaging with Axially Swept Light Sheet Microscopy. Biophys. J. 108, 2807. 10.1016/j.bpj.2015.05.013.

30. Syeda, R., Xu, J., Dubin, A.E., Coste, B., Mathur, J., Huynh, T., Matzen, J., Lao, J., Tully, D.C., Engels, I.H., et al. (2015). Chemical activation of the mechanotransduction channel Piezo1. Elife 4. 10.7554/eLife.07369.

31. Hedde, P.N., and Gratton, E. (2018). Selective plane illumination microscopy with a light sheet of uniform thickness formed by an electrically tunable lens. Microsc. Res. Tech. 81, 924–928. 10.1002/jemt.22707.

32. Power, G., Ferreira-Santos, L., Martinez-Lemus, L.A., and Padilla, J. (2024). Integrating molecular and cellular components of endothelial shear stress mechanotransduction. Am. J. Physiol. Heart Circ. Physiol. 327, H989–H1003. 10.1152/ajpheart.00431.2024.

33. Heiser, C.N., Wang, V.M., Chen, B., Hughey, J.J., and Lau, K.S. (2021). Automated quality control and cell identification of droplet-based single-cell data using dropkick. Genome Res. 31, 1742–1752. 10.1101/gr.271908.120.

34. Fu, H., Yu, Y., Wang, S., Xu, P., Sun, Y., Li, J., Ge, X., and Pan, S. (2025). Piezo1 disrupts blood–brain barrier via CaMKII/Nrf2 in ischemic stroke. Cell. Mol. Life Sci. 82, 259. 10.1007/s00018-025-05804-8.

35. Young, M.D., and Behjati, S. (2020). SoupX removes ambient RNA contamination from droplet-based single-cell RNA sequencing data. Gigascience 9, 1–10. 10.1093/gigascience/giaa151.

36. Harraz, O.F., Klug, N.R., Senatore, A.J., Hill-Eubanks, D.C., and Nelson, M.T. (2022). Piezo1 Is a Mechanosensor Channel in Central Nervous System Capillaries. Circ. Res. 130, 1531–1546. 10.1161/CIRCRESAHA.122.320827.

37. Germain, P.L., Robinson, M.D., Lun, A., Garcia Meixide, C., and Macnair, W. (2022). Doublet identification in single-cell sequencing data using scDblFinder. F1000Res. 10. 10.12688/f1000research.73600.2.

38. Jia, B.Z., Tang, X., Rossmann, M.P., Zon, L.I., Engert, F., and Cohen, A.E. (2025). Swimming motions evoke Ca2+ events in vascular endothelial cells of larval zebrafish via mechanical activation of Piezo1. bioRxiv, 2025.02.05.636757. 10.1101/2025.02.05.636757.

39. Shipley, F.B., Dani, N., Xu, H., Deister, C., Cui, J., Head, J.P., Sadegh, C., Fame, R.M., Shannon, M.L., Flores, V.I., et al. (2020). Tracking Calcium Dynamics and Immune Surveillance at the Choroid Plexus Blood-Cerebrospinal Fluid Interface. Neuron 108, 623. 10.1016/J.NEURON.2020.08.024.

40. Korsunsky, I., Millard, N., Fan, J., Slowikowski, K., Zhang, F., Wei, K., Baglaenko, Y., Brenner, M., Loh, P. ru, and Raychaudhuri, S. (2019). Fast, sensitive and accurate integration of single-cell data with Harmony. Nature Methods 2019 16:12 16, 1289–1296. 10.1038/s41592-019-0619-0.

41. Lim, X.R., Willemse, L., and Harraz, O.F. (2025). Amyloid beta Aβ1-40 activates Piezo1 channels in brain capillary endothelial cells. Biophys. J. 124. 10.1016/j.bpj.2024.12.025.

42. Gu, Z., Eils, R., and Schlesner, M. (2016). Complex heatmaps reveal patterns and correlations in multidimensional genomic data. Bioinformatics 32, 2847–2849. 10.1093/bioinformatics/btw313.

43. Kirst, C., Skriabine, S., Vieites-Prado, A., Topilko, T., Bertin, P., Gerschenfeld, G., Verny, F., Topilko, P., Michalski, N., Tessier-Lavigne, M., et al. (2020). Mapping the Fine-Scale Organization and Plasticity of the Brain Vasculature. Cell 180, 780–795.e25. 10.1016/j.cell.2020.01.028.

44. Yu, G., Wang, L.G., Han, Y., and He, Q.Y. (2012). clusterProfiler: an R Package for Comparing Biological Themes Among Gene Clusters. https://home.liebertpub.com/omi 16, 284–287. 10.1089/omi.2011.0118.

45. Perin, P., Rossetti, R., Ricci, C., Cossellu, D., Lazzarini, S., Bethge, P., Voigt, F.F., Helmchen, F., Batti, L., Gantar, I., et al. (2021). 3D Reconstruction of the Clarified Rat Hindbrain Choroid Plexus. Front. Cell Dev. Biol. 9, 692617. 10.3389/fcell.2021.692617.

46. Isaacs, D., Xiang, L., Hariharan, A., and Longden, T.A. (2024). KATP channel–dependent electrical signaling links capillary pericytes to arterioles during neurovascular coupling. Proc. Natl. Acad. Sci. U. S. A. 121, e2405965121. 10.1073/pnas.2405965121.

47. Stamenkovic, S., Li, Y., Waters, J., and Shih, A. (2023). Deep Imaging to Dissect Microvascular Contributions to White Matter Degeneration in Rodent Models of Dementia. Stroke 54, 1403–1415. 10.1161/STROKEAHA.122.037156.

48. Krolak, T., Kaplan, L., Navas, K., Chen, L., Birmingham, A., Ryvkin, D., Izsa, V., Powell, M., Wu, Z., Deverman, B.E., et al. (2025). Brain Endothelial Gap Junction Coupling Enables Rapid Vasodilation Propagation During Neurovascular Coupling. Cell 188, 5003. 10.1016/j.cell.2025.06.030.

49. Chuntharpursat-Bon, E., Povstyan, O. V., Ludlow, M.J., Carrier, D.J., Debant, M., Shi, J., Gaunt, H.J., Bauer, C.C., Curd, A., Simon Futers, T., et al. (2023). PIEZO1 and PECAM1 interact at cell-cell junctions and partner in endothelial force sensing. Communications Biology 2023 6:1 6, 358-. 10.1038/s42003-023-04706-4.

50. Császár, E., Lénárt, N., Cserép, C., Környei, Z., Fekete, R., Pósfai, B., Balázsfi, D., Hangya, B., Schwarcz, A.D., Szabadits, E., et al. (2022). Microglia modulate blood flow, neurovascular coupling, and hypoperfusion via purinergic actions. Journal of Experimental Medicine 219. 10.1084/jem.20211071.

51. Conway, D.E., and Schwartz, M.A. (2015). Mechanotransduction of shear stress occurs through changes in VE-cadherin and PECAM-1 tension: implications for cell migration. Cell Adh. Migr. 9, 335–339. 10.4161/19336918.2014.968498.

